# *Drosophila* Mutants that Are Motile but Respond Poorly to All Stimuli Tested: Mutants in RNA Splicing and RNA Helicase, Mutants in The Boss

**DOI:** 10.1101/066423

**Authors:** Lar L. Vang, Julius Adler

## Abstract

Adult *Drosophila melanogaster* fruit flies were placed into one end of a tube near to repellents (benzaldehyde and heat) and away from the other end containing attractants (light and a favored temperature). They escaped from the repellents and went to the attractants. Five motile mutants that failed to do that were isolated. They did not respond to any external attractants tested or external repellents tested. In addition, they did not respond well to internal sensory stimuli like hunger, thirst, and sleep. The mutants, although motile, failed to respond to stimuli at both 34°C and at room temperature. Some of the mutants have been mapped. The mutants are missing RNA splicing and RNA helicase. In addition, mutants missing information from The Boss are discussed.

## I. INTRODUCTION

Organisms are constantly exposed to a variety of external attractants and external repellents as well as to a variety of internal sensory stimuli. How organisms respond to these to bring about behavior is a basic question of life.

One approach for discovering how this works is the isolation and study of mutants that fail here. In this report we show that *Drosophila* flies can be mutated in such a way that, although still motile, they no longer respond well to any sensory stimulus tested. This includes various external attractants and various external repellents as well as internal sensory functions like hunger, thirst, and sleep. An account of some of this work has appeared (Lar Vang and Julius Adler, 2016, 2018). A preliminary report of some of the results has been presented (Adler, 2011; Vang, Alexei Medvedev, and Adler, 2012).

An unusual feature here is that a single mutation causes a great many different defects in behavior. This makes it likely that the cause is a substance with broad properties. It is now proposed that these mutants are defective in RNA splicing and RNA helicase.

Do organisms have a director? Another case where a single mutation would cause a great many different defects in behavior is “The Boss”. The idea of The Boss was introduced by Adler in “My Life with Nature”, p. 60 of Annual Review of Biochemistry, 2011: “The Boss is the thing inside every organism – humans, other animals, plants, microorganisms – that is in charge of the organism.” All things that an organism does are controlled by The Boss: The Boss controls behavior, metabolism, development, immunological response, and reproduction (Figure 10 of Adler, 2016). While so far The Boss has been just an idea, this idea may now be supported by mutants studied here: these mutant may fail to respond to sensory stimuli due to lacking the behavior part of The Boss.

Figure 1 summarizes a current view of the mechanism of behavior:

**Figure 1.**
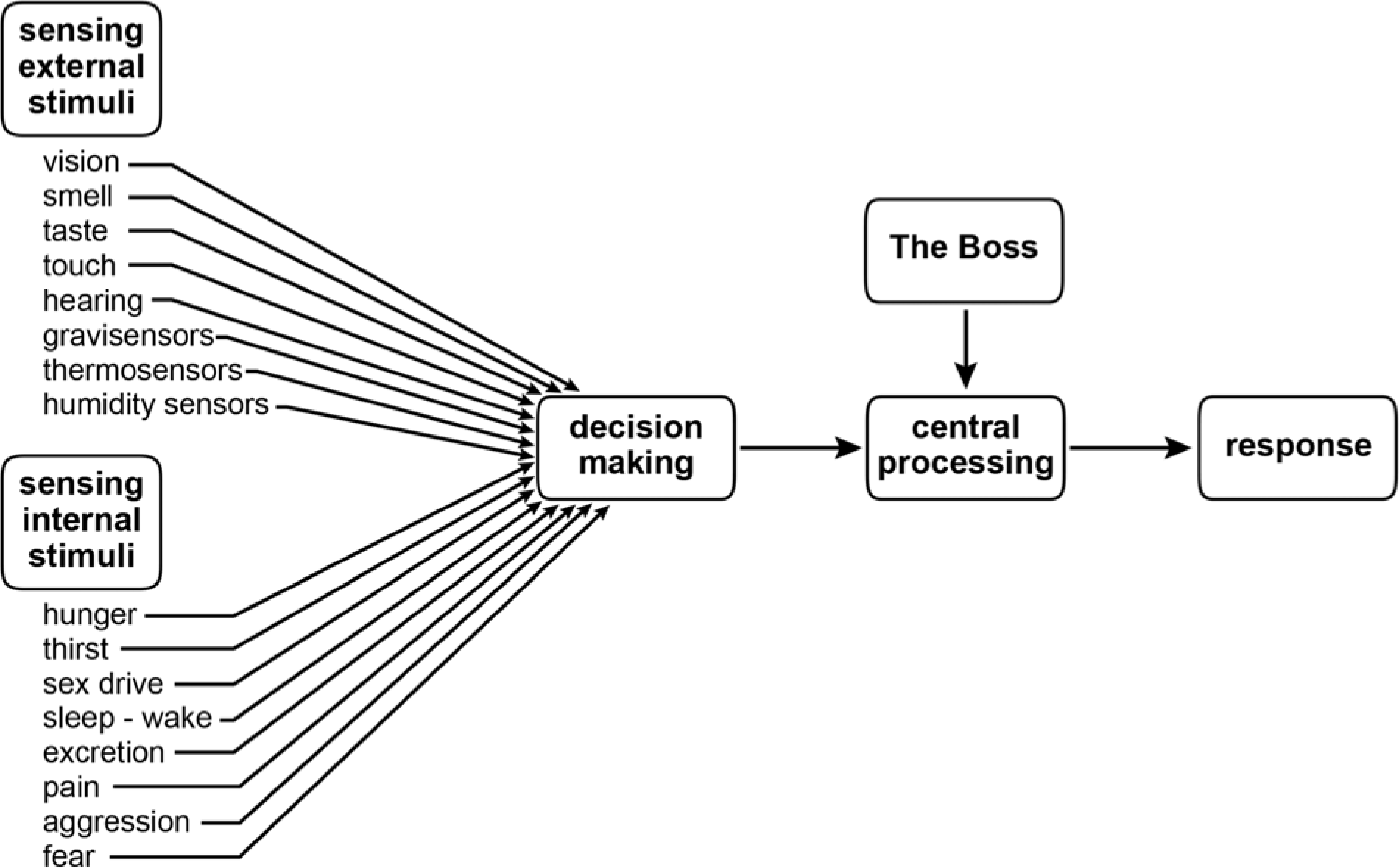
The mechanism of behavior. This applies to all organisms: microorganisms, plants, animals including humans. Decision making occurs when several sensory stimuli are encountered together, which is usually the case. Central processing includes behavior, metabolism, development, immunological response, and reproduction.

Work by others has shown that behavior in insects includes vision (Peter Weir and Michael Dickinson, 2015), smell and taste (Leslie Voshall and Reinhard Stocker, 2007), audition (Jan Clemens, Cyrille Girardin, Philip Coen and coworkers, 2015), and courtship (Haria Pavlou and Stephen Goodwin, 2013). Central processing includes the central complex, which is a system of neuropils consisting of the protocerebral bridge, the fan-shaped body, the ellipsoid body, and noduli (Ulrike Hanesch, Karl-Friedrich Fischbach, and Martin Heisenberg, 1989; Roland Strauss and Heisenberg, 1993; J.M. Young and Douglas Armstrong, 2010; Tanya Wolf, Nirmala Iyer, and Gerald Rubin, 2015). The relation of the central complex to the work reported here is described at the end.

## II. RESULTS

### A. RESPONSES TO EXTERNAL STIMULI

#### 1. RESPONSE TO STIMULI USED TOGETHER

In a 34°C dark room flies were started near two repellents (0.1M benzaldehyde and 37°C) at one end of a tube, away from two attractants (light at 1000 lux and 27°C) at the other end (Figure 2). The parent responded by going away from the repellents and to the attractants (Figure 3A). Mutants that were not motile were rejected, only the motile mutants were studied. This consisted of five mutants, named 1 to 5. Such a mutant, Mutant 2, failed to respond when the four stimuli were present (Figure 3B). Each of the four other mutants also failed to respond when the four stimuli were present (Vang and Adler, 2016).

**Figure 2.**
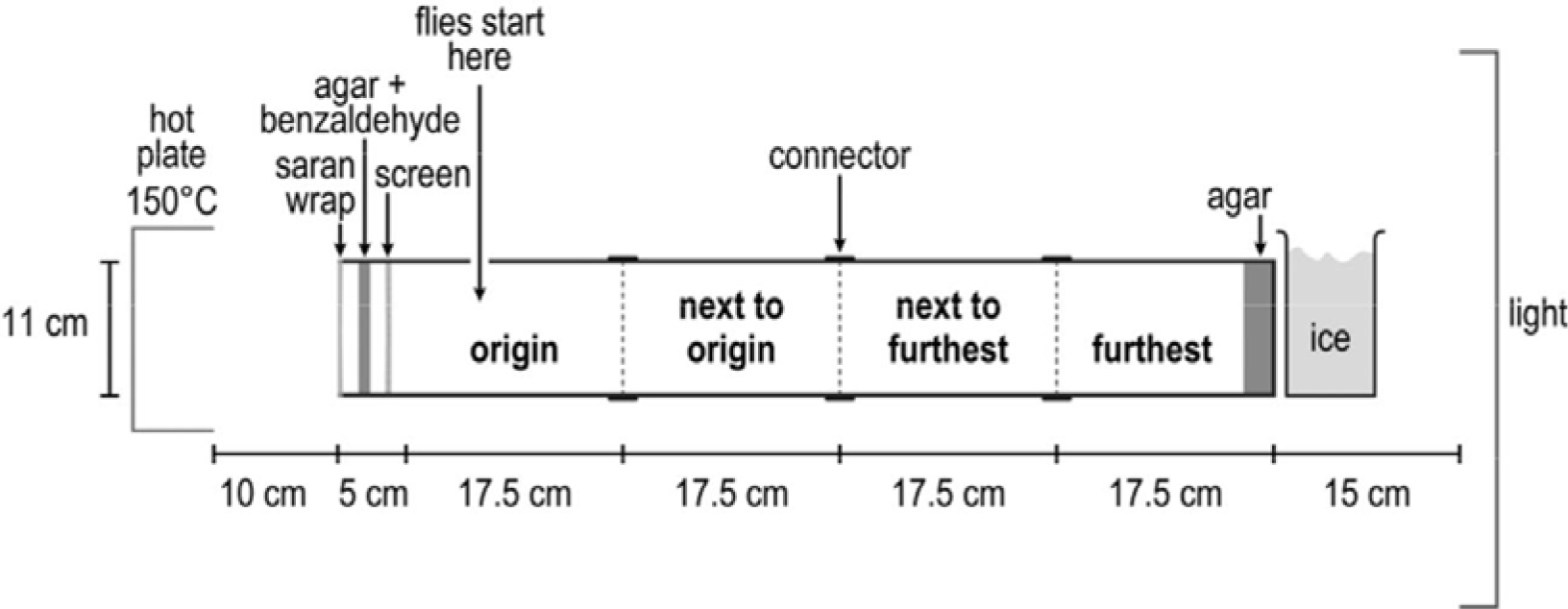
Apparatus for isolating and testing mutants in a 34°C room. At the left end were repulsive 0.1M benzaldehyde and repulsive 37°C (due to a hot plate at 150°C). At the right end were attractive light (1000 lux) and attractive 27°C (due to ice water). The middle was close to 34°C.

**Figure 3.**
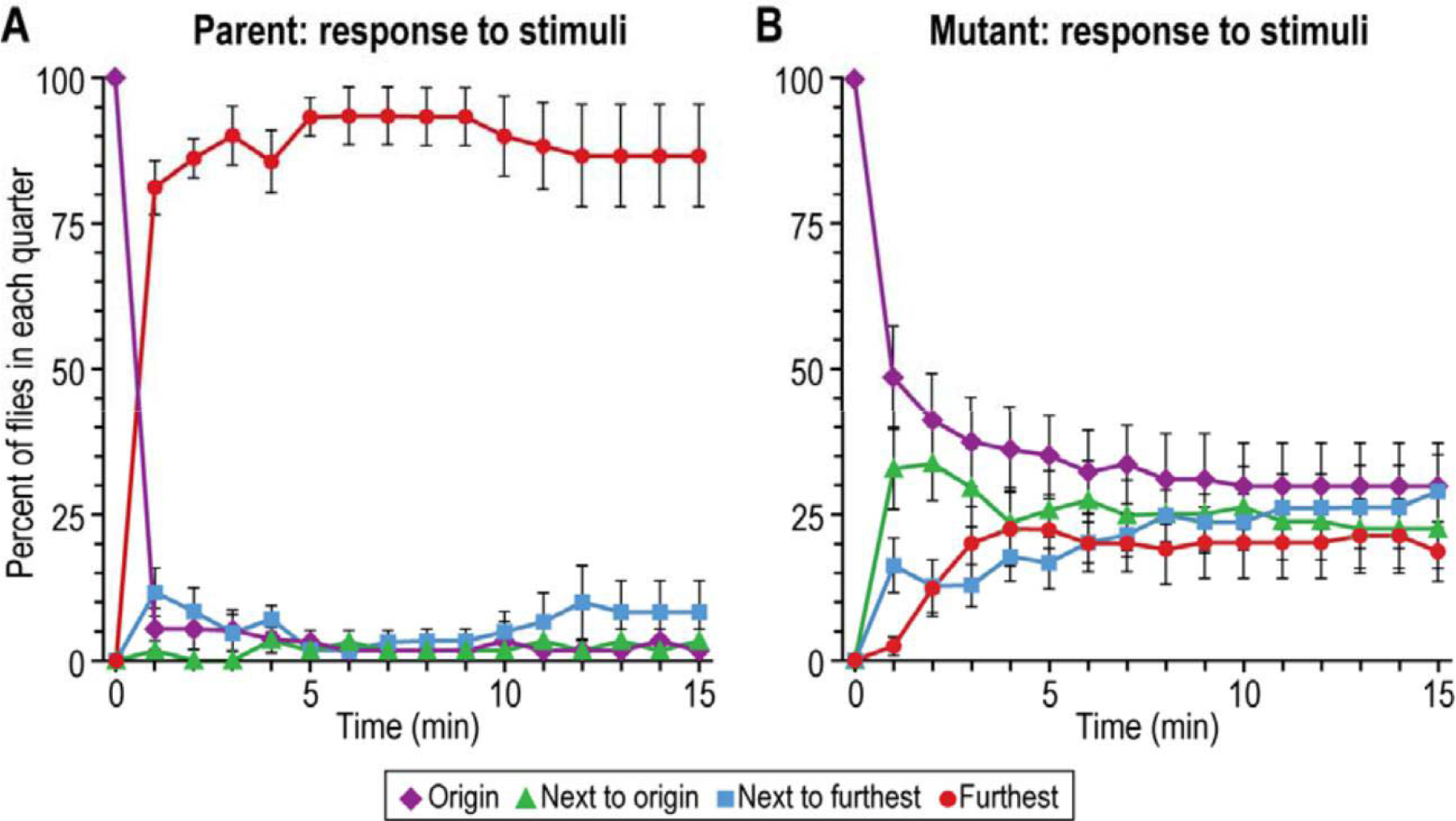
Response to stimuli used together. Repellents (0.1M benzaldehyde and high temperature (37°C) were at the left end, attractants (light, 1000 lux, and a favored temperature (27°C) at the right end. ***A***, Parental response (n=7). ***B***, Mutant 2 (n=8). Flies were tested in a 34°C room with 10 to 20 flies used per trial. Data are mean±SEM.

#### 2. RESPONSE TO INDIVIDUAL STIMULI

A single stimulus was presented to flies that were derived from ones that had already experienced the four stimuli. For example, the parent went to light only (Figure 4A) while a mutant did not (Figure 4B). Each of the five mutants failed to respond to light only (Vang and Adler, 2016).

**Figure 4.**
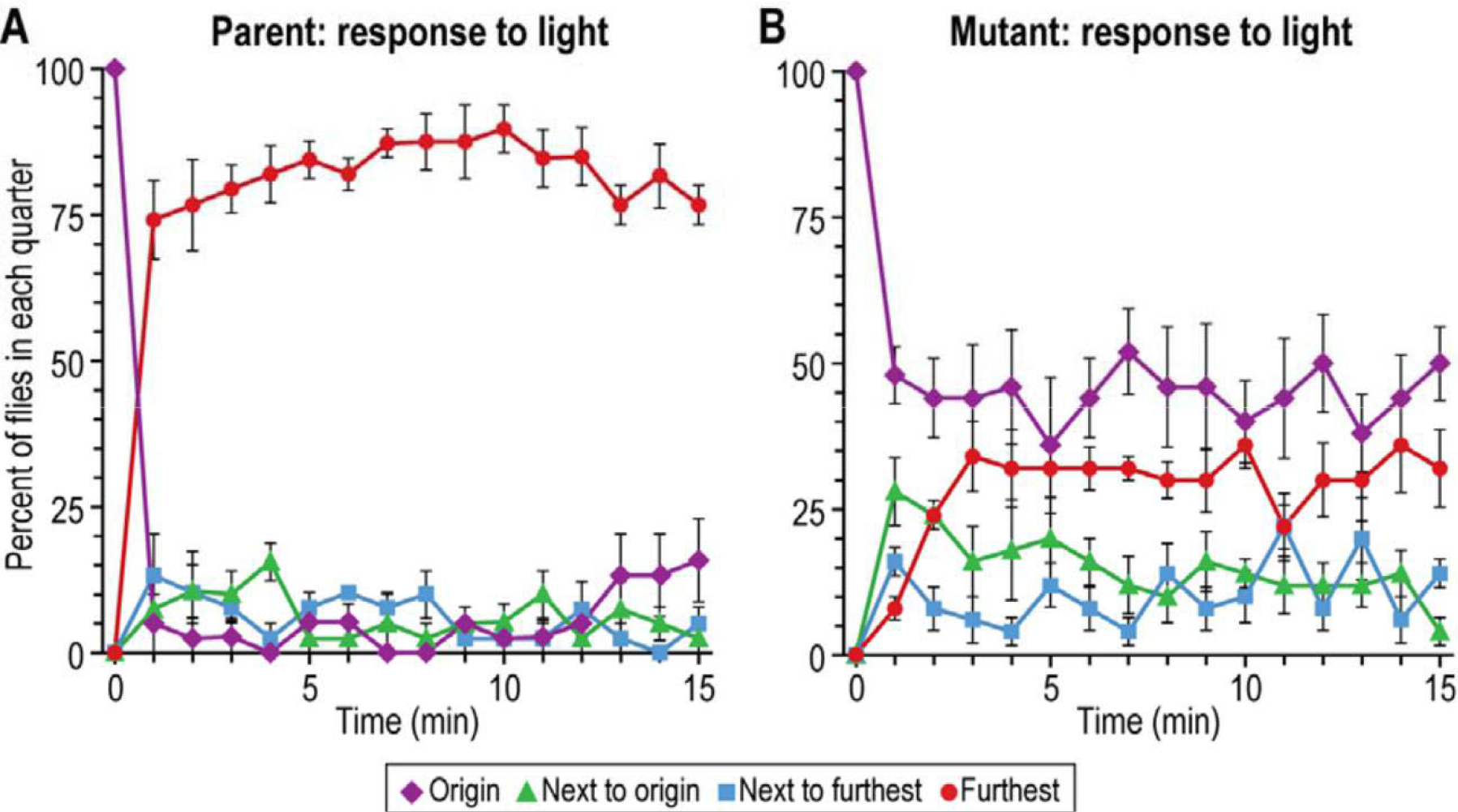
Response to light alone. Light (1000 lux) was placed at the right end as in Fig. 2. ***A***, Parental response (n=4). ***B***, Mutant 1 response (n=5). Flies were tested at 34°C with 10 to 20 flies used per trial. Data are mean±SEM.

For heat alone, the parent was repelled (Figure 5A) but the mutant was not repelled (Figure 5B). That was the case for Mutants 1 and 2 (Vang and Adler, 2016). (The other mutants were not tested for this.)

**Figure 5.**
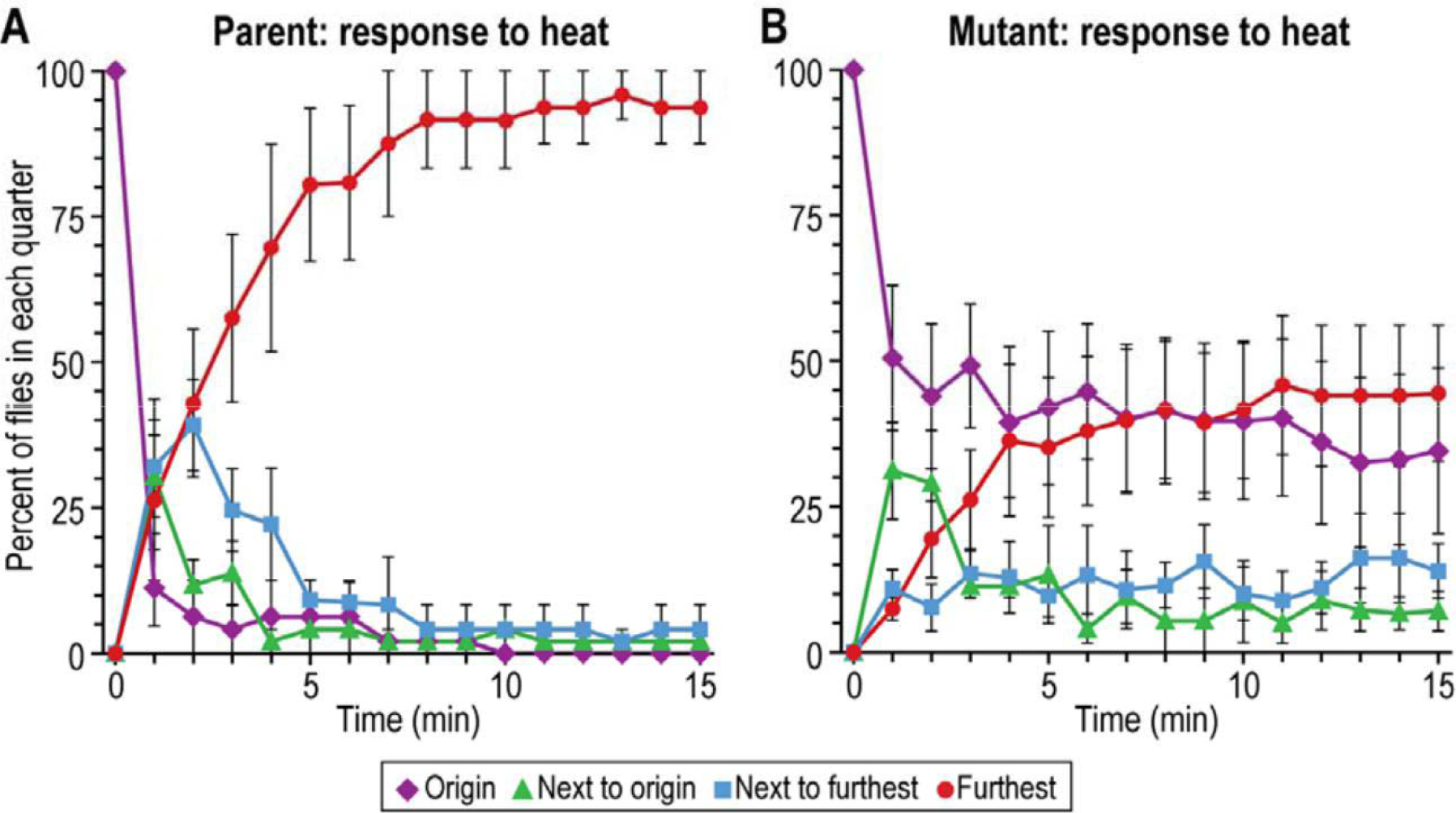
Response to heat gradient alone. The heat source was placed at the left end as in Figure 2. ***A***, Parental response (n=4). ***B***, Mutant 1 response (n=5). Flies were tested at 34°C with 10 to 20 flies per trial. The warm side measured 37°C and the cool side 27°C. Data are mean±SEM.

A similar result was found for benzaldehyde alone: the parent was repelled by benzaldehyde while Mutants 1 and 2 were not repelled. See Vang and Adler, 2016, for the figures. (The other mutants were not tested for this.)

Thus the mutants were defective not only for the four stimuli used together but also for each stimulus used alone.

#### 3. RESPONSE TO OTHER EXTERNAL STIMULI

These mutants were in addition tested with stimuli that were not among those four used to obtain the mutants:

The mutants were tested for response to the attractant sucrose after starvation (Robert Edgecomb, Cara Harth, and Anne Schneiderman, 1994) for 17 to 20 hours. Compared to the wild-type, both Mutants 1 and 2 consumed less sucrose, about 20% as much as the wild-type. See Vang and Adler, 2016, for the figures. (The other mutants were not tested for this.)

In the case of the repellent quinine, flies were started in a 0.1M quinine half and then they had the opportunity to go into a non-quinine half (see Vang, Medvedev, and Adler, 2012, for details of the method). The parent went into the non-quinine half but Mutant 1 and Mutant 2 did not. See Vang and Adler, 2016, for the figures. (The other mutants were not tested for this.)

To test response to gravity, these flies were placed into a vertical tube and pounded down, then at every minute the flies in each third of the tube were counted (see Vang, Medvedev, and Adler, 2012, for details of the method). The parent responded by climbing up while Mutants 1 and 2 climbed up 10% as well after subtraction of movement without any added stimuli. See Vang and Adler, 2016, for the figures. (The other mutants were not tested for this.)

Thus these mutants, isolated by use of the four stimuli, were defective even for stimuli that were not present during their isolation.

#### 4. MOVEMENT WITHOUT ANY ADDED STIMULI

In the absence of any stimulus added by the experimenters, the parent (Figure 6A) and the mutant (Figure 6B) moved similarly, indicating that motility alone is about the same in parent and mutant. This was found also for Mutants 2, 3, 4 and 5 (Vang and Adler, 2016). Aside from our seeing the flies, these results tell that the mutants are motile.

**Figure 6.**
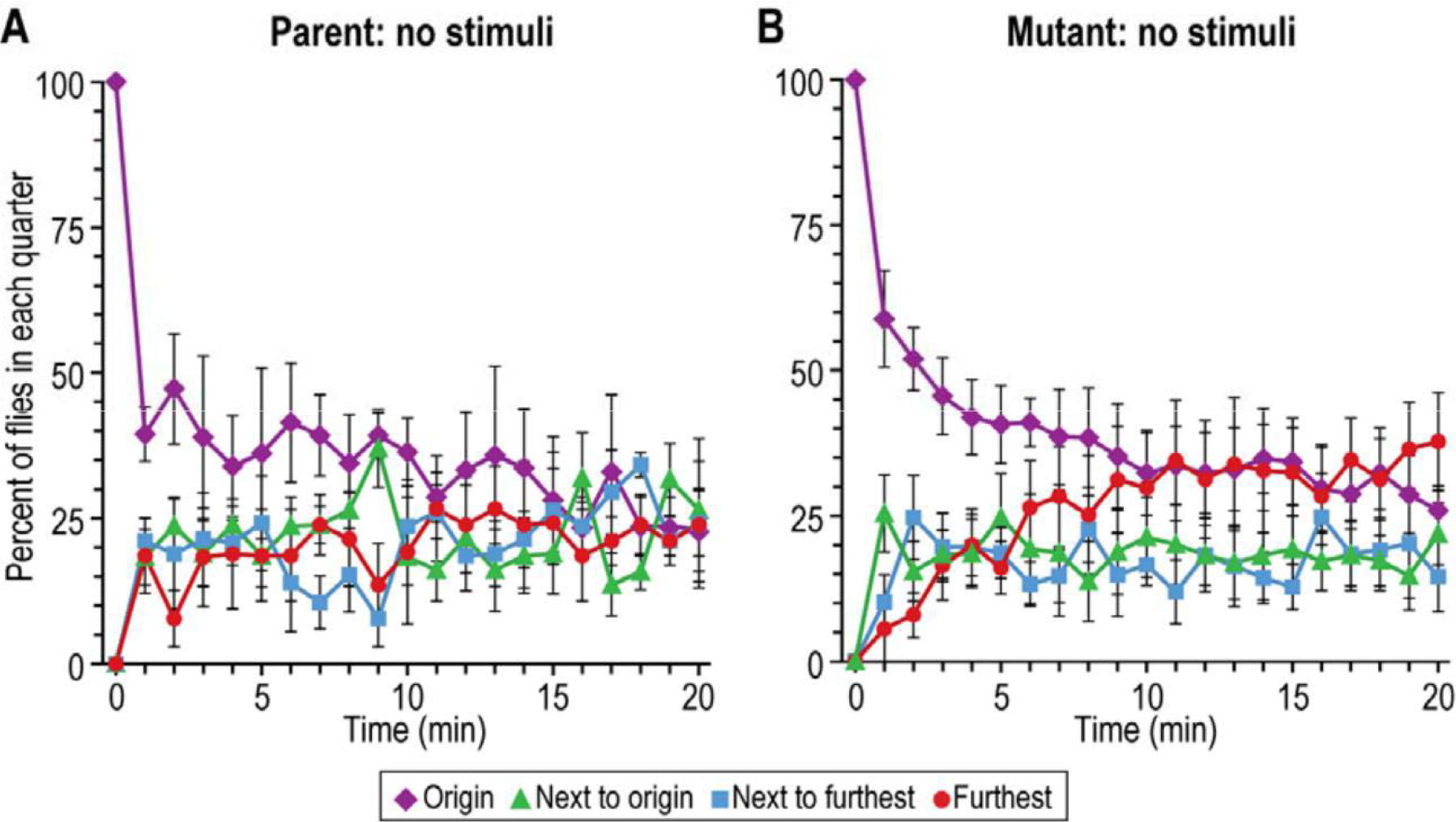
Response without added stimuli. ***A***, Parental response (n=4). ***B***, Mutant 1 response (n=6). Flies were tested at 34°C with 10 to 20 flies used per trial. Data are mean±SEM.

#### 5. EFFECT OF INCUBATION TEMPERATURE

All the work reported above was carried out in a 34°C room in order to allow, if necessary, isolation and study of conditional mutants, i.e. mutants defective at 34°C but not defective at room temperature. We measured response to light (1000 lux) at room temperature (21 to 23°C). The parent responded to light but all five of the mutants failed to respond to light or responded only 10% as well as the parent, just as they did at 34°C (Vang and Adler, 2016). Thus the mutations are not conditional.

Presumably these mutants are defective to all stimuli at room temperature, not just to light. Figures below show defects at room temperature for hunger, thirst, and sleep. Then how could the mutants survive and grow at room temperature? It must be that the mechanism studied here is not an essential one: flies live and reproduce without it.

### B. RESPONSES TO INTERNAL STIMULI

#### 1. HUNGER

Here we focus on hunger (Robert Edgecomb, C.E.Harth, and A. M. Schneiderman, 1994; Christoph Melche, Ruediger Barder, and Michael Pankratz, 2007; Katzuyo Fujikawa, Aya Takahashi, Asuza Nishimura and coauthors., 2009; Shelli Farhadian, Mayte Suarez-Farinas, Leslie Voshall and coauthors, 2012; Seung-Hyun Hong, Kyu-Sun Lee, Su-Jin Kwak and coauthors, 2012; Pavel Itskov and Carlos Ribeiro, 2012). To measure hunger we used an apparatus (Fig. 7), inspired and modied from an earlier design (L. Barton-Brown and D.R. Evans, 1960), that we described (Vang and Adler, 2016).

**Figure 7.**
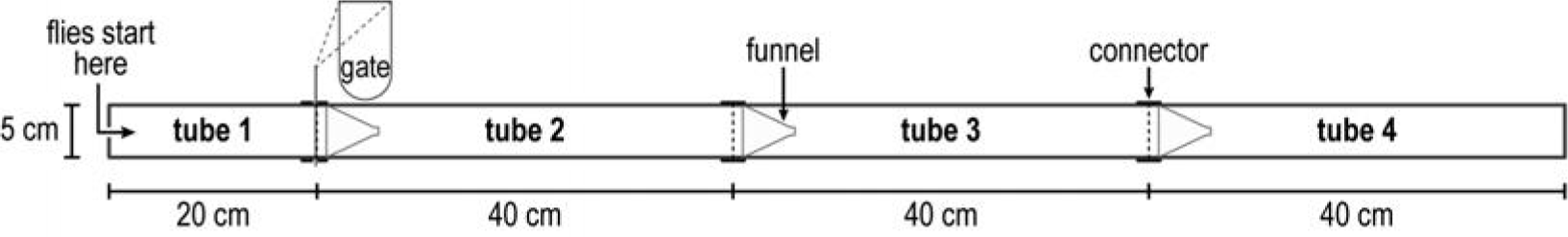
Apparatus for measuring hunger and for measuring thirst. For details see (Vang and Adler, 2016). Tube 1 is called “origin”. Flies were tested at room temperature (21-23°C) for up to 40 hours.

Briefly, in a dark room at 21-23°C male flies – parent or mutants - were transferred into one end (tube 1) of a 5 × 140 cm apparatus containing throughout its length a 5 cm wide strip of wet paper to satisfy thirst but containing no food. Starvation for food began once the flies were put in. Every 10 hours the location of the flies was measured with light on for a few seconds.

At 20 hours the parent had largely left the origin (tube 1) and had begun to accumulate at the end (tube 4) (Figure 8A, solid bars), while the mutant had moved towards the end very little (Figure 8B, solid bars). This is interpreted to mean that the parent is searching for food while the mutant is defective in searching for food.

**Figure 8.**
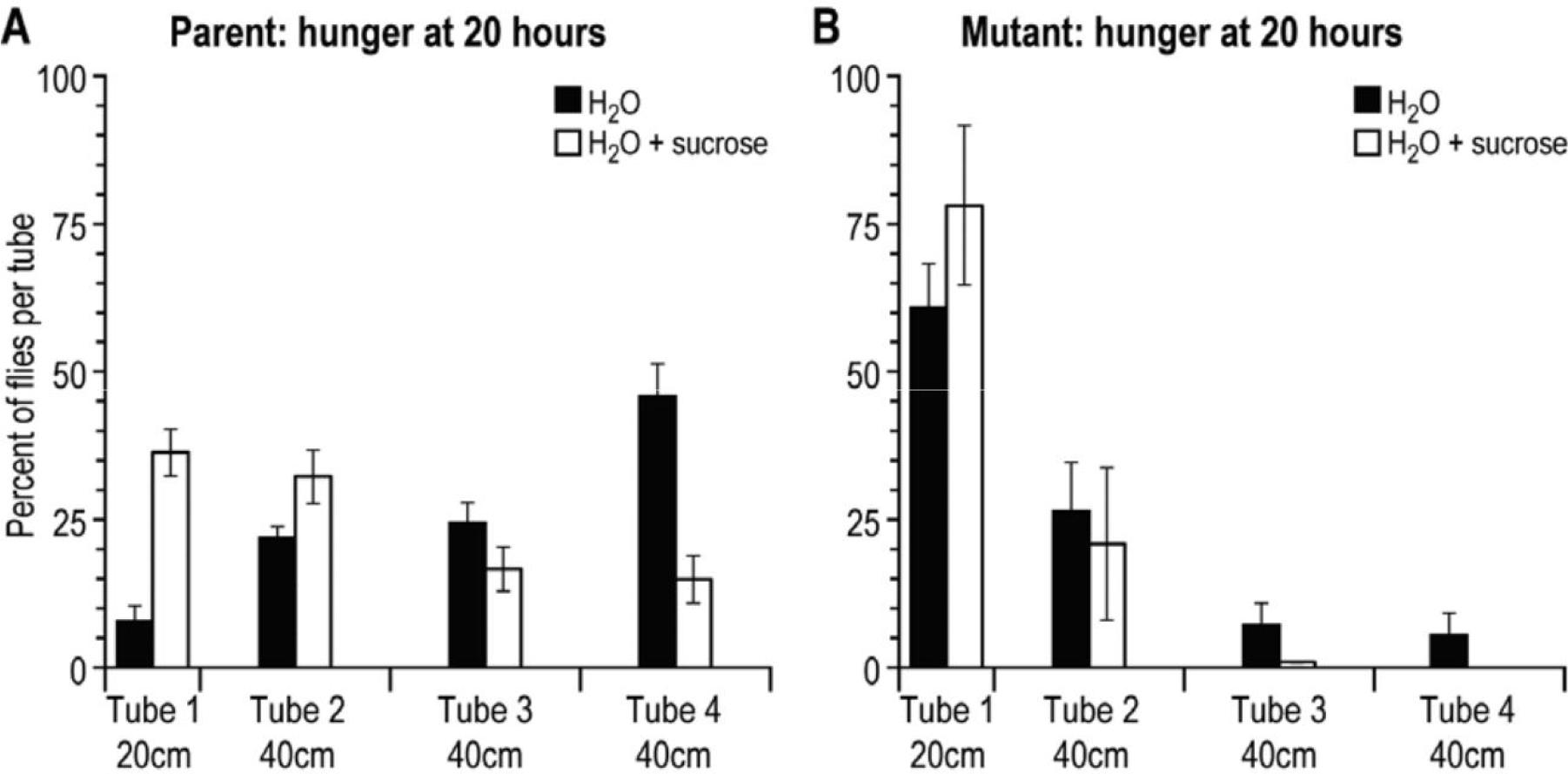
Movement of flies at 20 hours in search for food. Solid: water but no food (no sucrose). Open: water and food (0.1M sucrose). ***A***, Parental response with water only (n=5) and with water + sucrose (n=9). ***B***, Mutant 2 response with water only (n=5) and with water + sucrose (n=4). Data are mean±SEM. See (Vang and Adler, 2016) for Mutant 1; the other mutants were not tested for this. Flies were tested at room temperature (21-23°C) with 40 to 60 flies used per trial.

When food (0.1M sucrose) was added throughout the tube along with the wet strip of paper, the parent moved less far (rather than accumulating at the end) (Figure 8A open bars), while the mutant remained mostly where placed (Figure 8B, open bars). Since sucrose inhibited the movement of the parent, it is supposed that movement without sucrose is due largely to hunger. From these results we conclude that the mutants are defective in hunger.

#### 2. THIRST

To study thirst, flies were deprived of water. The procedure is the same as for hunger except that water was omitted and solid sucrose was layered throughout (Vang and Adler, 2016). Mutants 1 and 2 were tested, the other mutants not (Vang and Adler, 2016). By 30 hours the parent had moved out, presumably to search for water since addition of water inhibited this (Vang and Adler, 2016). The mutant moved out less well than the parent (Vang and Adler, 2016), so we conclude that the mutants are defective in thirst.

#### 3. SLEEP-WAKE

The parent and mutants isolated here were studied for sleep and wake according to the procedure of Cory Pfeiffenberger *et al.*, 2010. The parent was different from the mutants (Figure 9). The parent showed greatest activity at the start and end of the day but not in the middle of the day. Mutant 2 showed high activity throughout the day. Mutant 1 was less active than the parent at the start of the day.

**Figure 9.**
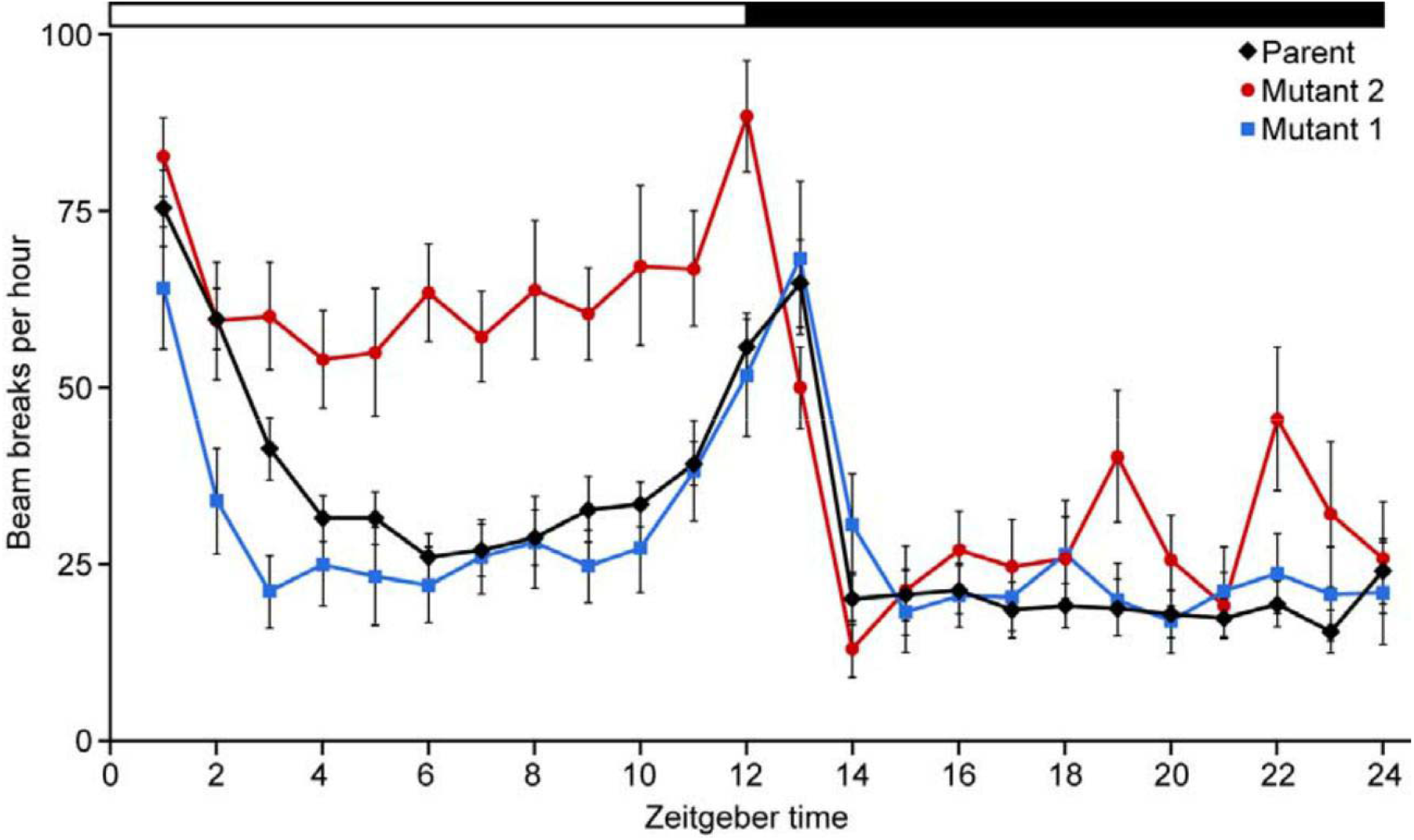
Circadian response. Individual flies are placed into a tube (5 × 20 mm) with an infrared light beam intersecting at the middle of the tube. Mutant 1 (n=24), Mutant 2 (n=24), and parental response (n=24) are recorded over a 24 hour period at 22° C. Data are mean±SEM. There are smaller differences between the parent and the three other mutants (Vang and Adler, 2016).

### C. MAPPING OF THE MUTANTS

We found that Mutant 1 maps in a small gap between 12E3 and 12E5 on the X-chromosome (Vang and Adler, 2016). According to “Gene: I(1)G0007 *D. melanogaster*” 12E3-12E5 is involved in RNA splicing and RNA helicase.

We found that Mutant 2 maps next to or in the *CG1791* gene, a part of the fibrinogen gene of the X-chromosome. (Vang and Adler, 2016). Mutant 2 might map in a gene for The Boss. Mutants 3, 4, and 5 were not mapped; perhaps they map in genes for The Boss.

## III. DISCUSSION

Here we describe the isolation and some properties of *Drosophila* mutants that are motile but yet they each fail in response to all external attractants and all external repellents tested (Figures 3-5) and also they are deficient in response to internal sensory stimuli tested (Figures 8 and 9). Thus, although the mutants are motile, they have:

decreased responsiveness to light
decreased responsiveness to heat and to favorable temperature
decreased responsiveness to repulsive chemicals (like benzaldehyde)
decreased responsiveness to sweet tastants (like sucrose)
decreased responsiveness to bitter tastants (like quinine)
decreased responsiveness to gravity
decreased responsiveness to hunger
decreased responsiveness to thirst
abnormality in some sleep

Because all of these different behaviors are defective in the mutants, it seems reasonable to say that there is a single place that is responsible, rather than a defect in each of the many different sensory receptors. So here is a place that when mutated leads to a defect in many places. What is this place? This newly discovered place is in genes for RNA splicing and RNA helicase (see Mapping of the mutants, above). Defects in these leads to loss of all behavior. RNA splicing removes the introns from pre mRNA to produce the final set of instructions for the synthesis of proteins. RNA helicases are enzymes that separate lagging and leading strands of an RNA helix using energy derived from ATP hydrolysis.

There may be another mechanism that, when mutated, leads to loss of all behavior. This is called “The Boss” (see Mapping of the mutants, above). It is proposed that The Boss controls responses to all sensory stimuli. See below.

The path from sensory stimuli to behavioral responses in vertebrates and flies has been compared by Nicholas Strausfeld and Frank Hirth (2013); they reported that the basal ganglia of mammals (the frontal cortex) and the central complex of insects (the superior medial protocerebrum) have multiple similarities and they suggest deep homology between these cerebral cortices, see Figure 10. Origin of the cerebral cortex has been studied by use of an ancient ancestor by Raju Tomer, Dedlev Arendt and coauthors (2010) and Heather Marlow and Arendt (2014).

**Figure 10.**
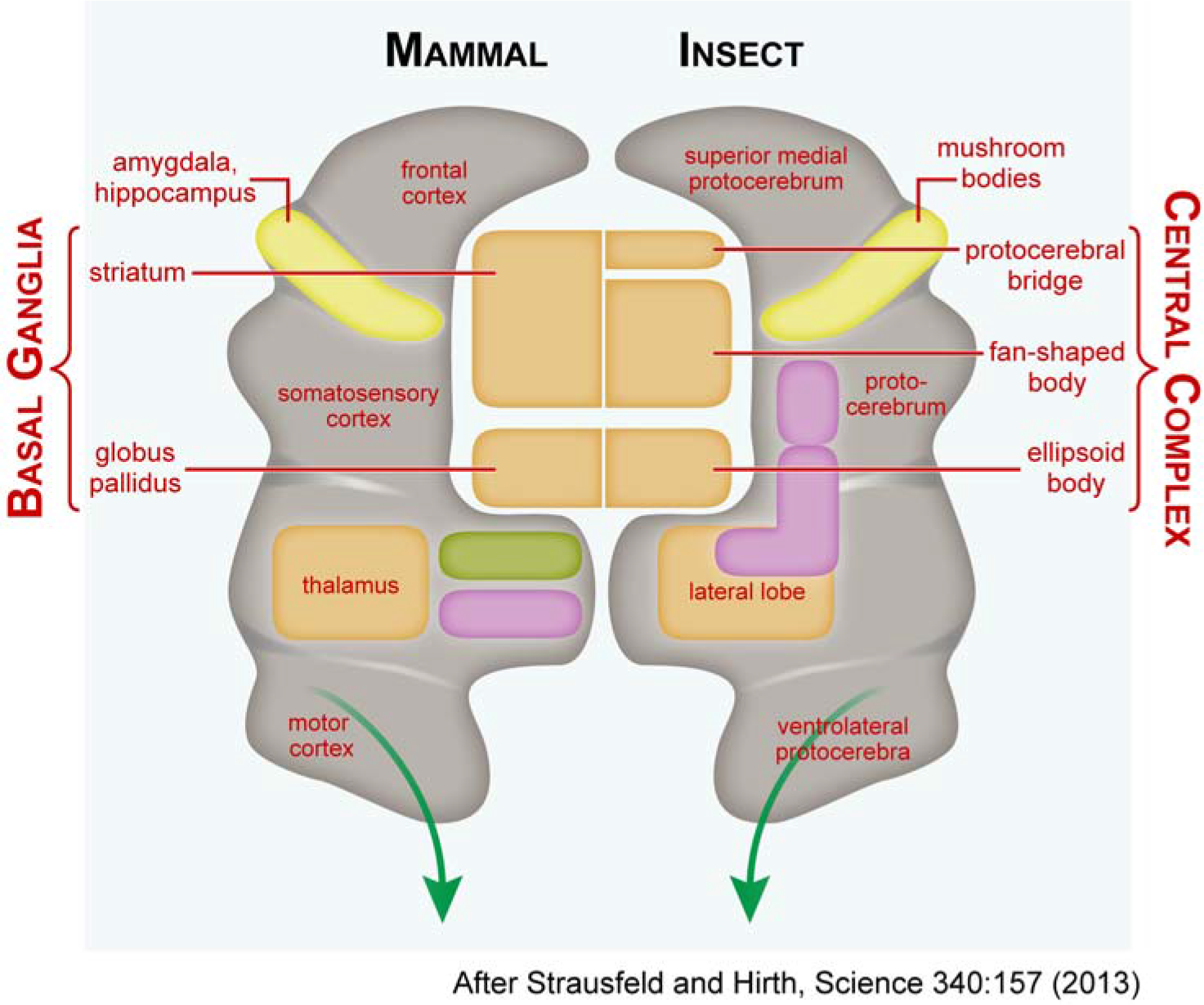
Comparison of the basal ganglia of vertebrates and the central complex of insects according to modification of Fig. 2 of Strausfeld and Hirth (2013). The green arrows lead to the response.

Starting in the 1870’s it became apparent to some psychologists that there is a part of the mammalian brain, the frontal cortex, that is master of the whole brain; see reviews up to 1970 by the neurophysiologist Aleksandr Luria, who himself modernized this concept and studied syndromes resulting from deficiencies of the frontal cortex (Luria, 1973, 1980). This part of the brain became known as the “central executive” through the research of the psychologist Alan Baddeley (1966) or as the “executive brain” through the research of the neuropsychologist Elkhonon Goldberg (2001), a student of Luria’s. It is now known as “executive control” or “executive function” as well as “prefrontal cortex”. For reviews see Goldberg and Dmitri Bougakov (2007), Joaquin Fuster (2008), Sam Gilbert and Paul Burgess (2008), Alfredo Ardila (2008), and Hyun Chung, Lisa Weyandt, and Anthony Swentosky (2014).

Helen Barbas reported (2000, 2013), as described in Figure 11, that the orbitofrontal cortex of humans and other primates receives information from the sensory cortices (which contain visual, auditory, somatosensory, gustatory, and olfactory data as it is received) and also from the amygdala (which contains data about memory and emotion) and also from the thalamus. Barbas suggested that “the orbitofrontal cortex is thus capable of sampling the entire external and internal environment and may act as an environmental integrator” (2000). The orbitofrontal cortex is a part of the prefrontal cortex (p. 3 of Barbas, 2013). After receiving all this information, the prefrontal cortex decides what action should be brought about by informing the premotor cortex (Barbas, 2013), whose function is to produce action. The prefrontal cortex has been reported also in rodents (Verity Brown and Eric Bowman, 2002; Harry Uylings, Henk Groenwegen, and Bryan Kolb, 2003). For a review of what constitutes the prefrontal cortex in mammals, rats, and mice see Marie Carlen (2017).

**Figure 11.**
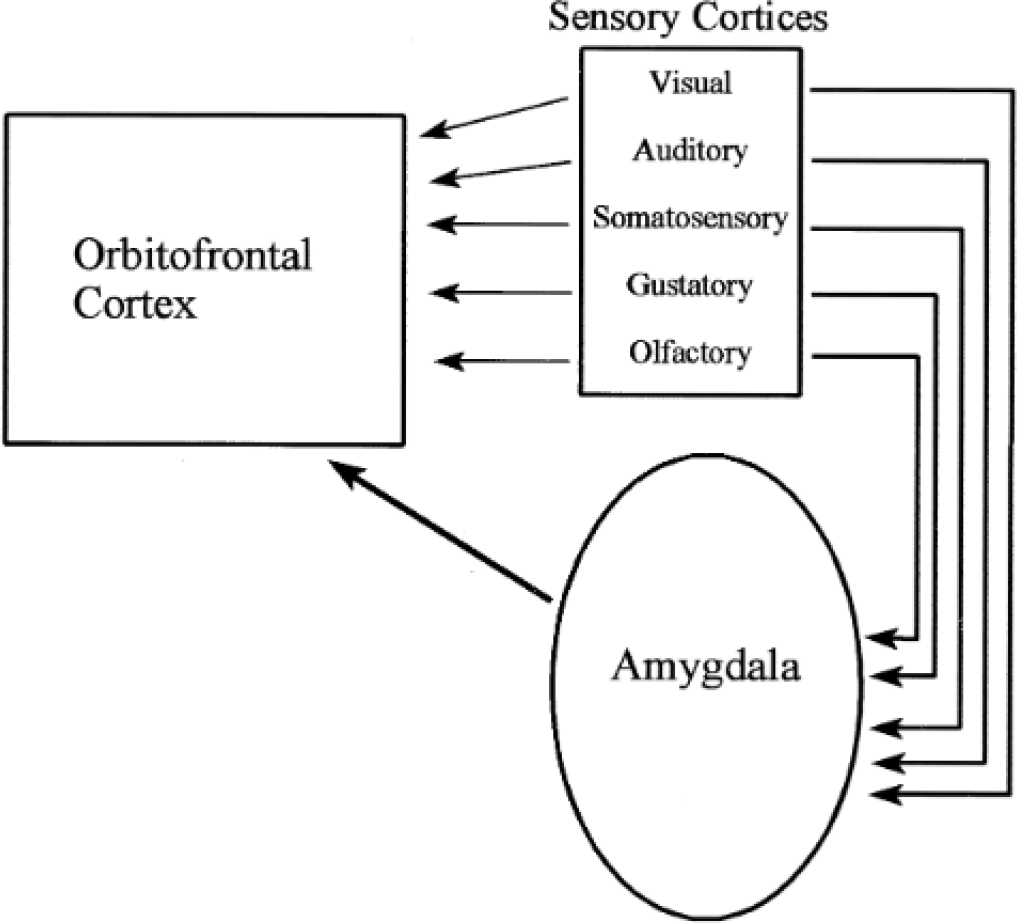
The orbitofrontal cortex receives information from sensory cortices and also from the amygdala’s recording of past events and emotions and also (not shown here) from the thalamus, as described in Fig. 3 of Barbas (2000). For more detail, see Figs. 41.14 and 41.31 of Barbas (2013).

It is suggested by us that the excitatory pathways and the inhibitory pathways of the prefrontal cortex (Figure 41.14 of Barbas, 2013) are directed by what is here called the “The Boss”. The Boss would be in charge of the prefrontal cortex, The Boss would control the prefrontal cortex. The Boss can turn the prefrontal cortex on or off depending for example o wake or sleep. We proposed that The Boss is to be found throughout biology, in animals, plants, and microorganisms, and that The Boss directs each organism (Adler, 2011, “My life with nature”, page 60; Adler, 2016, “A search for The Boss: The thing inside each organism that is in charge”). Thus every organism has something that is in control of the organism, namely The Boss.

The prefrontal cortex of vertebrates occurs in the frontal cortex (Barbas, 2013), which corresponds to the superior medial protocerebrum of insects: see Figure 10 above. The frontal cortex and the superior medial protocerebrum might well have analogous functions, so a system like that in the prefrontal cortex might occur in insects, too. The anatomy of the superior medial protocerebrum has been studied in flies, see James Phillips-Portillo and Strausfeld, 2012, and Cynthia Hsu and Vikas Bhandawat, 2016. The superior medial protocerebrum could respond to The Boss.

The control of genes by The Boss could be studied by finding conditional mutants in The Boss: these could be missing the response to all stimuli at some higher temperature but would be normal at room temperature in order to keep The Boss intact there. In the present report such mutants did not show up, the mutants described here failed at 34°C but they also failed at room temperature. However, only five mutants were studied here; possibly if a much larger number of mutants were isolated there would be some among them that are defective at 34°C but normally responsive at room temperature. In our work on decision making (Adler and Vang, 2016) mutants were isolated that fail at 34°C but do respond normally at room temperature, see next.

Decision making (see Figure 1) has been studied in animals, plants, and microorganisms. Decision making occurs in primates and rodents, as described by Jacob Cloke, Derek Jacklin, and Boyer Winters (2015). For a review of decision making in mice, *Drosophila*, and *Caenorhabditis elegans* see Nilay Yapici, Manuel Zimmer, and Ana Domingos (2014); about *Drosophila* they say, “Nutrient sensing that guides decision-making has also been studied in *Drosophila* and found to be regulated at the central nervous system level…These neurons are located in the lateral protocerebrum”. According to Shiming Tang and Aike Guo (2001) in *Drosophila* integration of conflicting sensory information (repellent heat of one color plus non-repellent room-temperature of another color) takes place in the mushroom body. According to Laurence Lewis, Gerald Rubin, Ilona Grunwald Kadow and coworkers (2015) in *Drosophila* integration of conflicting sensory information (repellent CO_2_ plus attractant vinegar) requires the mushroom body. *Drosophila* mutants were isolated by Adler and Vang (2016) that fail in making a decision (repellent methyl eugenol plus attractant light) at 34°C but the mutants respond normally in making a decision at room temperature, so it seems that The Boss could be in control of decision making and central processing. These *Drosophila* mutants (Adler and Vang, 2016) should be studied further to find out if some of them may act in the superior medial protocerebrum, see above, to bring about the central processing mentioned in Figure 1.

In conclusion for a portion here: The Boss is in control of the organism – behavior, metabolism, development, immunologic response, and reproduction – and defective behavior in mutants of The Boss is discussed here. The Boss activates the central complex, which is studied by use of mutants with defects in the different genes of the central cortex (Strauss and Heisenberg, 1993; Adler and Vang, Fig. S15, 2016).

## V. METHODS

Details of methods used here are found in in the previous paper (Vang and Adler, 2016): A. Isolation of mutants. B. How to study response to external stimuli. C. How to study response to internal stimuli.

## ACKNOWLEDGEMENTS

Julius Adler is grateful to the Camille and Henry Dreyfus Foundation for six years of grants in support of 31 undergraduate research students named in Adler and Vang, 2016, and in Vang and Adler, 2016. Lar Vang is currently associate research specialist in the Adler laboratory. Robert A. Kreber, a research specialist in Barry Ganetzky’s laboratory, has helped us in studies of the genetics of our mutants. We are grateful to Erin Gonzales and Jerry Yin for showing us how to use the *Drosophila* activity monitory system. Julius Adler thanks Barry Ganetzky for teaching him about fruit flies. Thanks to Millard Susman for criticism. We are thankful to Laura Vanderploeg for the art work.

